# Vessel-Resolved Mapping of Perivascular Spaces Reveals Hierarchical Neurofluid Organization *In Vivo*

**DOI:** 10.64898/2026.05.01.722268

**Authors:** Xiaoqing Alice Zhou, Xiaochen Liu, Weitao Man, Sang Cheon Choi, David Hike, Xinyue Wang, Yuanyuan Jiang, Maiken Nedergaard, Kristopher T. Kahle, Brian Bacskai, Xin Yu

## Abstract

Perivascular spaces (PVS) are central to cerebrospinal fluid (CSF)-interstitial fluid (ISF) exchange in the brain, yet their brain-wide organization and in vivo accessibility remain poorly resolved due to limited spatial resolution and contrast specificity of existing imaging approaches. Prior gadolinium (Gd)-enhanced MRI studies demonstrate global tracer distribution but yield spatially diffuse signals that do not resolve PVS at the level of individual vessels. Here we introduce an ultra-high-resolution dual-contrast MRI framework that enables vessel-resolved mapping of PVS across the whole brain in vivo. This approach combines ultra-high-field imaging, an implantable radiofrequency coil for enhanced local sensitivity, and intraventricular Gd delivery to achieve sufficient contrast and spatial specificity for detecting vessel-associated perivascular signal. Using this framework, we show that PVS are hierarchically organized along vascular trees, extending from major surface arteries into deep cortical and subcortical regions. Signal patterns along arterial branches and junctions indicate that PVS follows vascular topology. Quantitative analysis reveals that only a subset of penetrating vessels (∼6%) exhibits detectable PVS signal, indicating heterogeneous organization across vascular networks independent of vessel caliber. Widespread detection of PVS, including in the hippocampus, further demonstrates that ventricularly delivered tracers access a distributed, vessel-associated perivascular network *in vivo*. These results establish an anatomical framework for mapping perivascular transport pathways across the brain, bridging global tracer imaging with vessel-resolved organization, and enabling investigation of how vascular architecture and fluid dynamics shape CSF-ISF exchange.

## Introduction

Perivascular spaces (PVS) are fluid-filled compartments surrounding cerebral vessels and constitute a central anatomical interface for cerebrospinal fluid (CSF) and interstitial fluid (ISF) exchange in the brain^1-3^. Within the glymphatic framework, periarterial pathways have been proposed to support CSF influx and solute transport^4^, whereas alternative models emphasize intramural periarterial drainage along vascular basement membranes^5-7^. Although the relative contributions of these pathways remain debated, there is broad consensus that perivascular compartments play a fundamental role in brain fluid transport, solute clearance, and homeostasis, with dysfunction implicated in aging, cerebrovascular disease, and neurodegeneration^8-10^.

Recent work further indicates that CSF-ISF exchange is dynamically regulated by neural, vascular, and autonomic processes rather than passive diffusion alone. Brain state strongly modulates glymphatic transport, with sleep and circadian rhythms influencing fluid exchange and clearance^11^. In parallel, increasing evidence demonstrates tight coupling between neuronal activity and CSF dynamics, In particular, coordinated neural oscillations and slow-wave activity can modulate, and in some cases directly drive, brain-wide CSF movement and clearance^12-14^.Vascular dynamics also contribute to this process, as arterial pulsations and vasomotion have been implicated in shaping perivascular fluid movement^15^, while recent work shows that slow norepinephrine-dependent vasomotion during sleep promotes glymphatic clearance^16^. Moreover, neuromodulatory systems, including cholinergic and adrenergic pathways, can regulate vascular tone and CSF flux, further linking neural activity to fluid transport^17,18^. Together, these findings support a model in which PVS-mediated transport emerges from coupled neurovascular and neurofluid interactions.

Despite these conceptual advances, the anatomical organization and *in vivo* accessibility of PVS across the brain remain poorly resolved. In humans, conventional MRI primarily detects enlarged PVS as macroscopic fluid-filled structures, and recent intrathecal gadolinium (Gd) studies have begun to visualize periarterial compartments around large vessels, but these approaches remain limited to coarse spatial scales and CSF-accessible regions^19-21^. In rodents, contrast-enhanced MRI has demonstrated brain-wide tracer transport following CSF delivery and provided important evidence for CSF-ISF exchange; however, the resulting signals are spatially diffuse and reflect system-level tracer distribution rather than the precise geometry of perivascular spaces surrounding individual vessels^22-25^. Optical imaging approaches offer micron-scale resolution but are inherently limited in penetration depth, field of view, and physiological perturbation due to tracer delivery and surgical preparation. As a result, there remains no imaging framework capable of mapping PVS globally while resolving their organization at the level of individual vessels across cortical, subcortical, and deep brain regions *in vivo*.

Here, we address this gap by developing an ultra-high-resolution *in vivo* MRI framework for vessel-resolved mapping of PVS across the whole brain. By integrating ultra-high-field imaging, an implantable radiofrequency coil for enhanced local sensitivity, and dual-contrast strategies combining vascular signal suppression with intraventricular Gd delivery, we achieve sufficient sensitivity and contrast-to-noise ratio to detect perivascular signal at the scale of individual vessels. This approach enables whole-brain imaging at 30 μm isotropic resolution and complementary single-vessel mapping at 20 × 20 μm^2^ in-plane resolution. Using this framework, we demonstrate that PVS are hierarchically organized along vascular trees, extending from major surface arteries into penetrating cortical vessels and deep subcortical structures, including the hippocampus. These results establish a previously unavailable anatomical framework for resolving brain-wide perivascular organization in vivo and provide a foundation for linking vascular architecture to CSF-ISF transport dynamics under physiological and pathological conditions.

## Results

### Ultra–high-resolution mapping reveals perivascular spaces (PVS) along major cerebral vessels

Using 14 T MRI, we achieved 30 μm isotropic *in vivo* whole-brain imaging in 77 minutes, a spatiotemporal regime not previously attainable with MRI, enabling brain-wide visualization of PVS along major cerebral arteries. By integrating iron-based susceptibility contrast to suppress intravascular signal with intraventricular Gd infusion to selectively enhance CSF compartments, we developed a dual-contrast mapping strategy that enables simultaneous delineation of vascular lumens and adjacent perivascular spaces (PVS) (Fig. 1A). Co-registration of inflow-based MR angiography with Gd-enhanced images revealed a spatial correspondence between hypointense vessel lumens and adjacent hyperintense perivascular signals along major branches of the anterior cerebral artery (ACA). At the level of individual vessels, high-magnification views demonstrated a clear separation between the vessel core and a surrounding ring-like enhancement consistent with PVS (Fig. 1B). Also, Gd-enhanced PVS signals were prominently observed at T-shaped branch points of the ACA-azygos vessels (e.g., Animal 2), suggesting Gd transportation to hierarchical perivascular compartments along major arterial bifurcations. The hierarchical PVS of middle cerebral artery (MCA) were well identified in the ultra-high-resolution dual-contrast images (Supplementary Movie 1).

**Figure 1.**
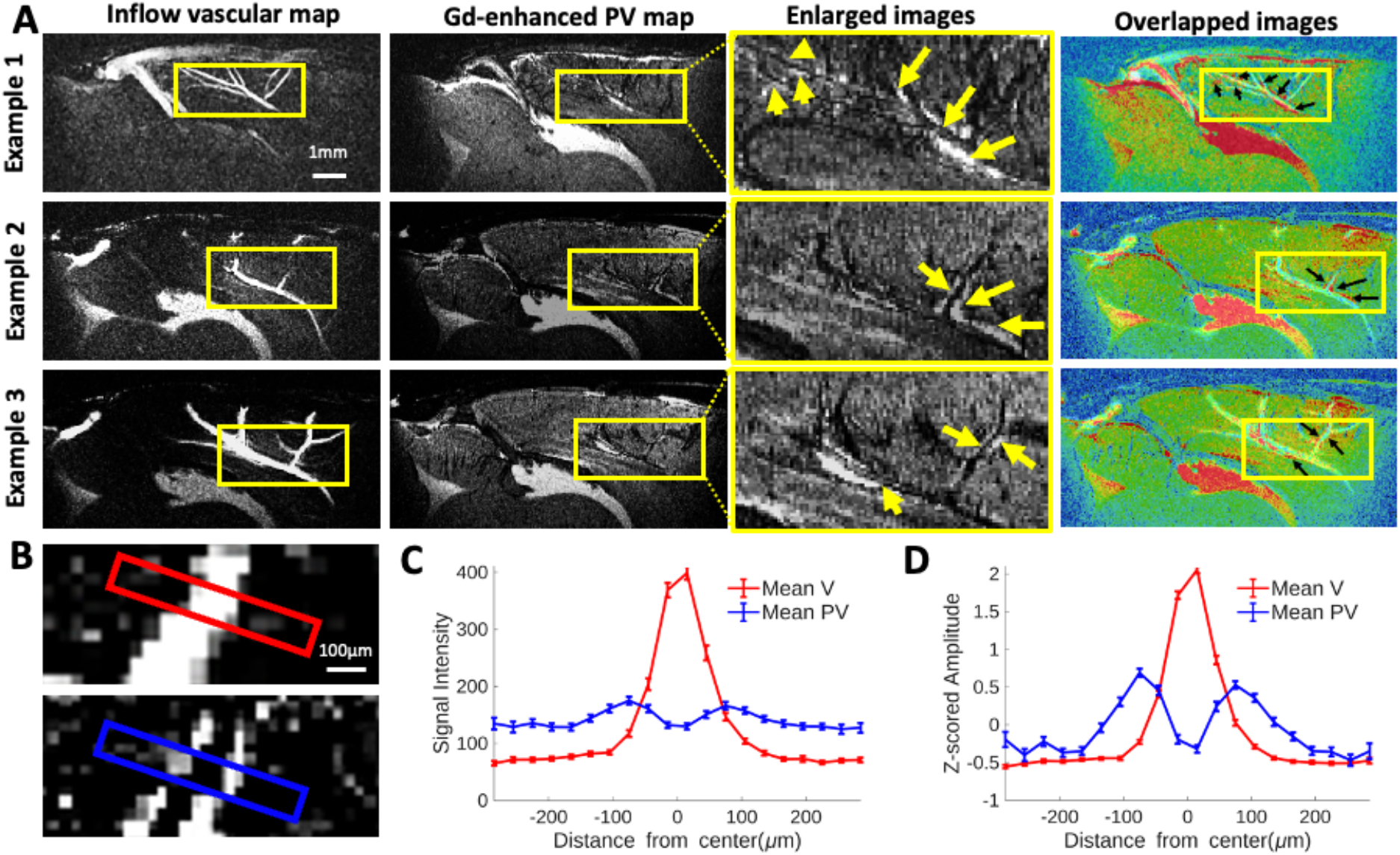
Ultra-high-resolution visualization of PVS along major cerebral arteries. **A**, Co-registered inflow-based vascular map (V), Gd-enhanced perivascular (PV) map, and their overlap demonstrate spatial correspondence between vessel lumens and adjacent perivascular spaces along major branches of the anterior cerebral artery (ACA) across animals (3 examples). Iron-based contrast suppresses intravascular signal, rendering vessels hypointense, while Gd infusion selectively enhances perivascular compartments (Enlarged images: arrows indicate Gd-enhanced PVS structure). **B**, Representative magnified view of a single ACA (azygos) vessel and its surrounding PVS. The top panel shows the inflow vascular image, and the bottom panel shows the corresponding Gd-enhanced PV image. Line profiles were extracted along the indicated axes (red and blue lines) over a 20-voxel segment (±10 voxels from vessel center; voxel size = 30 μm). **C**, Averaged intensity profiles across 150 vessels from 9 animals, showing spatial signal distribution from the vessel lumen (center) to surrounding perivascular regions. Profiles reveal a central hypointense vascular core flanked by enhanced perivascular signal. **D**, Normalized intensity profiles corresponding to (C), highlighting consistent spatial organization of vessel and perivascular signals independent of absolute intensity differences.

To quantitatively characterize the spatial relationship between PVS and vessel lumens, we extracted intensity profiles across vessels and their surrounding compartments. The resulting line profiles revealed a sharp central signal dip corresponding to the vessel lumen, flanked by pronounced peaks in the adjacent perivascular regions (Fig. 1C). After normalization, these profiles demonstrated a consistent spatial organization in which perivascular enhancement was confined to regions immediately adjacent to the vessel wall (Fig. 1D). Together, these findings established robust and spatially resolved visualization of PVS along major cerebral vessels at unprecedented *in vivo* resolution.

### Layer-resolved characterization of perivascular spaces in cortical penetrating vessels

We next asked whether the dual-contrast framework could resolve PVS associated with intracortical penetrating vessels, which constitute a critical interface for CSF-ISF exchange. Strikingly, high-resolution coronal imaging enabled continuous and unambiguous visualization of PVS along individual penetrating vessels across cortical depth, with perivascular signals tightly co-localized with vessel trajectories (Fig. 2A). This direct, slice-by-slice tracking reveals a coherent vessel-PVS architecture extending from the cortical surface into deeper layers, establishing clear delineation of perivascular compartments at the level of single penetrating vessels.

**Figure 2.**
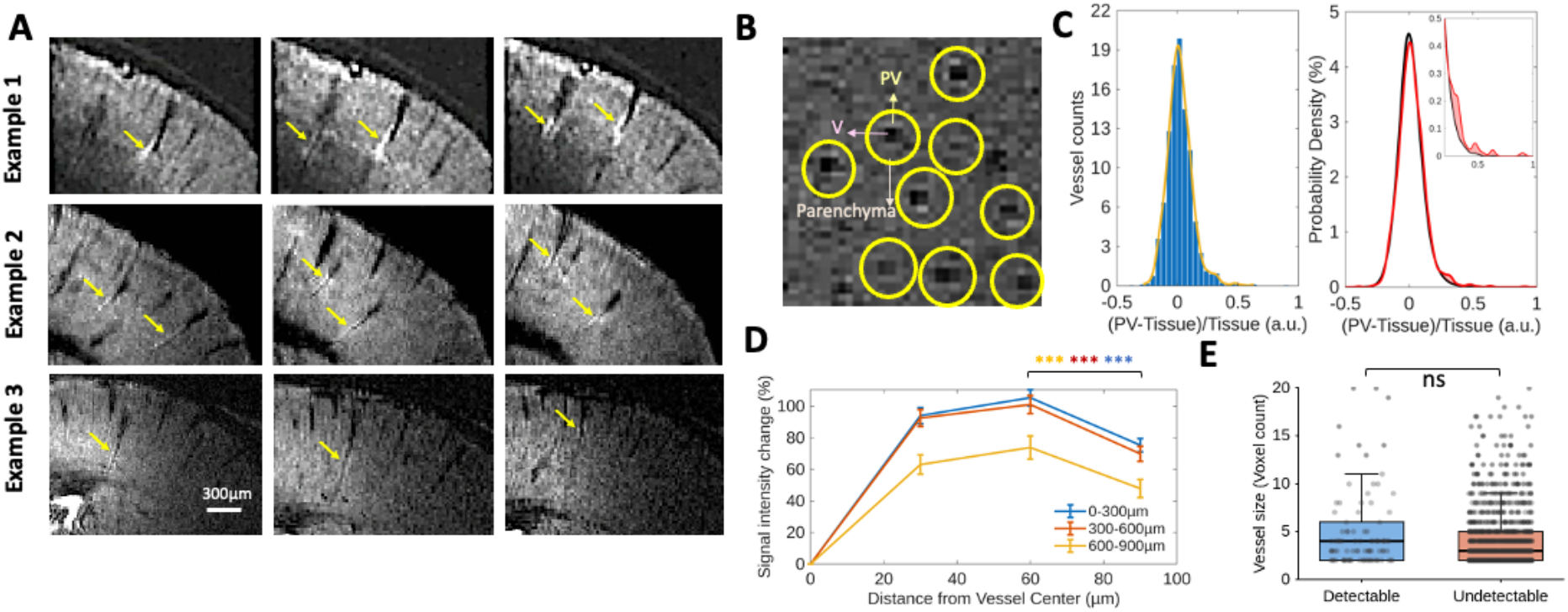
Layer-resolved characterization of PVS in cortical penetrating vessels. **A**, Representative coronal views showing penetrating cortical vessels and associated perivascular signals across animals. Arrows indicate vessel trajectories and adjacent PVS along cortical depth. **B**, Schematic of region-of-interest (ROI) selection and analysis workflow. The underlying image shows an axial view of cortical penetrating vessels. Vessel lumens were segmented using adaptive thresholding. Concentric perivascular layers were generated by iterative dilation (layer thickness: 30-60 μm), and surrounding parenchyma was defined as regions located 90 μm from the vessel boundary. **C**, Distribution of perivascular signal contrast across segmented vessels (n=1564 vessels from 8 animals). The left panel shows the empirical histogram of vessel-wise contrast values. The right panel shows the probability density of the observed distribution (red) together with a fitted normal distribution (black), highlighting deviation from normality and a right-skewed tail consistent with a subset of vessels exhibiting elevated perivascular signal. The inset shows a magnified view of the right tail of the distribution. D, Mean signal intensity profiles across vessel cross-sections derived from vessels with detectable PVS signal at three cortical depths (0–300, 300–600 and 600–900 μm from the pial surface; surface, middle and bottom layers). Signal profiles were sampled perpendicular to the vessel axis to obtain cross-sectional measurements. Data are presented as mean ± SEM (n = 237 vessels from 8 animals, p<0.001, FDR-corrected), showing elevated perivascular signal relative to the vessel lumen and surrounding parenchyma across depths. **E**, Comparison of vessel size between vessels with detectable and undetectable PVS signal. Vessel size does not differ significantly between the two groups, indicating that PVS detectability is not driven by vessel caliber. Data are presented as mean ± SEM (p=0.475). Each dot represents one vessel.

To enable systematic quantification, we developed an automated pipeline for vessel detection and perivascular segmentation. Vessel lumens were identified using adaptive thresholding to account for local intensity variations, and concentric perivascular layers were generated by iterative morphological dilation, defining radial compartments surrounding each vessel (Fig. 2B). Quantitative analysis was performed across multiple depths along penetrating vessels to ensure robust sampling of perivascular organization. The distribution of perivascular-to-tissue (PV/Tissue) signal ratios revealed a clear deviation from a randomized baseline. Fig. 2C shows that in contrast to the permutation-derived null Gaussian distribution, the observed vessel-based distribution exhibited a pronounced rightward shift, with an excess population in the high-ratio tail. There are approximately 6% of penetrating vessels contributing to this right-tail excess, exhibiting PV/Tissue ratios exceeding the range predicted by the null distribution. These vessels therefore represent a distinct subset with detectable perivascular enhancement. The absence of a comparable tail in the randomized distribution further supports that this effect reflects structured, vessel-associated signal rather than stochastic variation. Together, these results demonstrate that detectable perivascular tracer accumulation is restricted to a subset of penetrating vessels, revealing a heterogeneous and non-uniform organization of perivascular pathways across the cortical vasculature.

In addition, layer-resolved signal profiles of the selected penetrating vessels based on the histogram distribution demonstrated a consistent spatial gradient with signal intensity increased from the vessel lumen to inner perivascular layers, followed by a decrease toward surrounding parenchyma (Fig. 2D). The peak perivascular signal intensity exceeding adjacent tissue confirms the presence of detectable PVS compartments at the level of individual penetrating vessels. It should be noted that the size of the vessels showing strong PVS enhancements is not significantly different from the vessels without salient PVS enhancements (Fig. 2E), indicating that this PVS enhanced signals are vessel-size independent, further ruling out magnetic field disturbance-related artifacts. These findings establish that ultra-high-resolution MRI can resolve PVS not only along large surface vessels but also within cortical microvascular networks, with laminar specificity at tens-of-microns scale.

### Perivascular enhancement reflects CSF tracer accumulation rather than vascular or susceptibility artifacts

To determine whether the observed perivascular signals reflect CSF tracer accumulation rather than imaging artifacts, we performed controlled experiments that dissociate the effects of iron and Gd agents. During Gd infusion, robust perivascular enhancement was observed surrounding vessels. In contrast, images acquired 3 h post-infusion, after Gd washout but with iron remaining in the vasculature, retained pronounced vessel hypo-intensity while showing a marked loss of perivascular signal (Fig. 3A). Quantitative analysis of matched regions of interest confirmed this dissociation, demonstrating elevated perivascular signal during Gd infusion that returned to baseline following Gd clearance, despite persistent iron-induced signal suppression within vessel lumens (Fig. 3B,C). To further exclude local susceptibility as the source of perivascular enhancement, we compared phase maps with corresponding magnitude images at the level of individual vessels (Supplementary Fig. 2). There are some vessels exhibiting strong local field offsets, which reflected as pronounced phase shifts, but not showing corresponding perivascular signal enhancement. In contrast, vessels with robust perivascular signal displayed minimal phase perturbation. This direct dissociation indicates that perivascular enhancement is not coupled to local field inhomogeneity and cannot be explained by iron-induced susceptibility effects. Together, these results demonstrate that perivascular enhancement is not attributable to iron-induced susceptibility or flow-related artifacts, but instead reflects the accumulation of Gd-based CSF tracer within PVS compartments.

**Figure 3.**
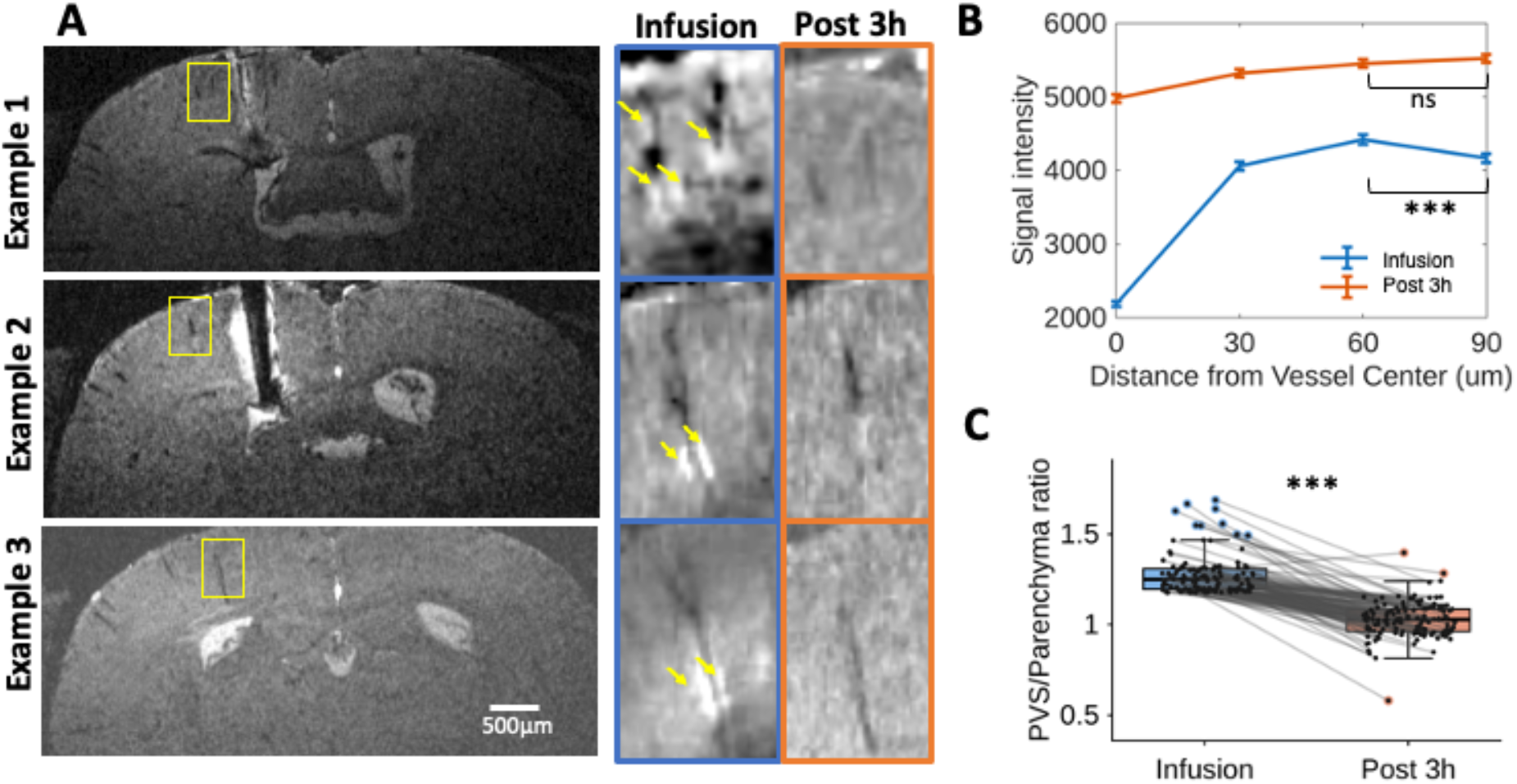
Perivascular enhancement arises from CSF tracer accumulation rather than susceptibility artifacts. **A**, Representative high-resolution MR images acquired during Gd infusion (left, blue) and 3 h post-infusion (right, orange) across animals. Yellow boxes indicate regions containing penetrating vessels. Vessel lumens remain hypointense in both conditions due to persistent iron contrast, whereas perivascular enhancement is observed only during Gd infusion and is absent after Gd washout. **B**, Mean signal intensity profiles across vessel lumen and concentric perivascular layers for the two conditions. Data are presented as mean ± SEM (n = 200 vessels from 8 animals). Perivascular signal is significantly higher during Gd infusion compared to 3 h post-infusion at matched distances from the vessel center (*P < 0.001, FDR-corrected), while vascular contrast remains unchanged, confirming that PVS enhancement reflects CSF tracer accumulation rather than iron-induced or flow-related artifacts. **C**, Perivascular-to-parenchyma signal ratio for individual vessels during Gd infusion and 3 h post-infusion. Data are presented as mean ± SEM (n = 200 vessels from 8 animals). Each dot represents one vessel. The PV/parenchyma ratio is significantly higher during infusion compared to post-infusion (*P < 0.001), and decreases following Gd clearance, indicating that perivascular enhancement reflects transient CSF tracer accumulation.

### Multi-scale dynamic mapping links ventricular CSF transport to hippocampal vessel-resolved PVS

To establish the spatiotemporal dynamics of tracer transport following intraventricular delivery, we performed time-resolved mapping of Gd distribution across ventricular and parenchymal compartments. Gd signal rapidly dispersed from the lateral ventricles into surrounding brain regions and persisted over an extended period (∼120 min), indicating sustained tracer retention and widespread CSF-ISF exchange (Supplementary Fig. 3, Supplementary Movie 2). These measurements define a mesoscale transport regime characterized by tracer entry from ventricular CSF into brain tissue, in particular, the hippocampal parenchyma, and gradual washout over ∼2 hours, establishing a well-defined temporal window during which tracer is present within parenchymal and perivascular compartments.

Within this defined transport window, we next investigated the spatial localization of tracer at the level of individual vessels using a 2D ultra-high-resolution single-vessel imaging scheme^26^ (in-plane resolution: 20 × 20 μm^2^) in the hippocampus. At baseline, hippocampal vessels were detected with hyperintense signals due to the blood inflow effect. Following intravenous iron infusion, blood signals within vessels were markedly attenuated by T2* effects, producing strong hypointensity that clearly delineates vessel lumens in single-vessel maps. Subsequent intraventricular Gd infusion introduced a distinct and spatially confined enhancement surrounding the vessels, indicating the Gd-enhanced PVS (Fig. 4). This dual-contrast modulation produced a characteristic configuration of dark vessel lumens encircled by bright perivascular rims (Fig. 4A,C), enabling direct separation of vascular and perivascular compartments at the level of individual vessels. Quantitative intensity profiles across inflow, iron, and Gd conditions further confirmed this compartmental dissociation, with peak signal localized to perivascular regions and suppressed within the vessel lumen (Fig. 4D). Together, these results established vessel-resolved detection of hippocampal PVS *in vivo*, linking ventricular CSF tracer transport to a brain-wide PVS network.

**Figure 4.**
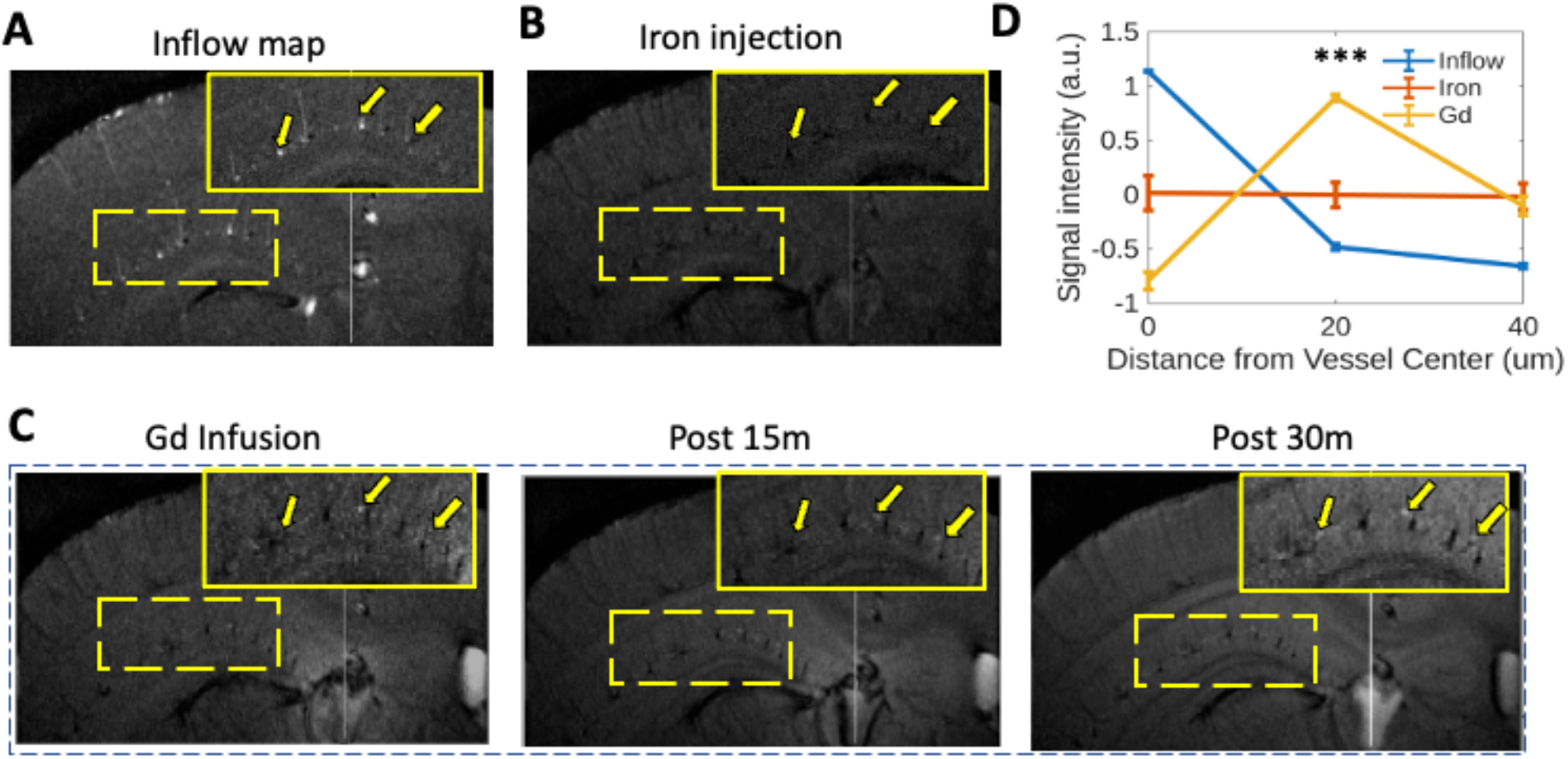
Dynamic contrast modulation separates vascular and perivascular signal components in penetrating vessels. **A-C**, Sequential MR images acquired at baseline (**A**), after iron infusion (**B**), and after Gd infusion (**C**) demonstrate distinct signal contributions from vascular and perivascular compartments. At baseline, vessels exhibit strong inflow signal; following iron infusion, vessels become hypointense due to T2* effects; subsequent Gd infusion produces localized enhancement surrounding the vessel, consistent with perivascular space labeling. Yellow boxes highlight representative regions. **D**, Mean signal intensity profiles across vessel-centered ROIs for the three conditions. Data are presented as mean ± SEM (n = 38 vessels from 8 animals). Iron selectively suppresses intravascular signal, whereas Gd selectively enhances perivascular compartments, enabling clear separation of vascular and perivascular signals. Perivascular signal under Gd is significantly higher than both inflow and iron conditions (P < 0.001, FDR-corrected).

## Discussion

In this study, we established an ultra-high-resolution, dual-contrast MRI framework that enables global, *in vivo* mapping of PVS surrounding individual vessels across the brain. Whole-brain imaging at 30 μm isotropic resolution enabled direct, vessel-resolved delineation of PVS across the brain *in vivo*. The integration of iron-based susceptibility contrast with intraventricular Gd infusion enabled direct separation of vessel lumens from adjacent PVS, delineating vessel-PVS organization across vascular hierarchies. This approach revealed continuous PVS along penetrating cortical vessels, extended to deep brain regions including the hippocampus, and was supported by complementary single-vessel imaging at 20 × 20 μm^2^ resolution. Importantly, controlled contrast dissociation experiments demonstrated that the observed perivascular signals reflected tracer accumulation rather than susceptibility- or flow-related artifacts. Together, these findings directly addressed a central unresolved question, whether PVS can be mapped globally *in vivo* at the level of individual vessels, and established a framework for vessel-resolved characterization of perivascular architecture across the brain.

### State of the Art and Limitations of MRI-Based PVS Imaging

Early Gd-enhanced MRI studies in rodents have established that tracers introduced into CSF can access the brain beyond the subarachnoid space, producing widespread parenchymal enhancement and providing initial *in vivo* evidence of CSF-ISF exchange^23-25^. Following cisterna magna infusion, contrast agents were observed to enter periarterial pathways along major surface arteries and propagate into deeper brain regions, enabling mapping of brain-wide tracer distribution^23^. Subsequent work using quantitative Gd-DOTA imaging and dynamic MRI further characterized the temporal evolution of tracer influx and clearance, demonstrating that a substantial fraction of delivered contrast enters and persists within brain parenchyma^24,25^. More recent studies have shown that these signals reflect global tracer transport dynamics modulated by vascular activity and neuromodulatory state, leading to spatially widespread and diffuse enhancement pattern^17^. Despite these advances, the resulting MRI signals primarily reflect system-level tracer distribution rather than the anatomical organization of PVS at the level of individual vessels. A recent in vivo Gd-enhanced MRI study mapped tracer transport along pial periarterial PVS, showing that these spaces can serve as conduits for convective CSF entry into brain tissue^27^. This approach primarily captured macroscopic pial PVS flow and downstream tracer distribution, rather than resolving the fine intracortical PVS architecture around penetrating vessels. Notably, emerging high-resolution *in vivo* MRI studies have begun to report vessel-associated signal features, including enhancement patterns localized near hippocampal vessels, suggesting potential sensitivity to perivascular compartments at finer spatial scales^22^. However, such signals remain difficult to interpret unambiguously, as prior studies have largely inferred their perivascular origin based on anatomical patterns and disease-related changes, without direct MRI-based suppression or separation of intravascular inflow contributions from hippocampal vessels^28^.

In humans, Gd-enhanced MRI has been used to characterize CSF-brain exchange at the whole-brain level, revealing spatially heterogeneous tracer accumulation and clearance over extended time scales^29^. Intrathecal imaging studies further demonstrate tracer propagation along vessel-associated pathways and enhancement in fluid-filled compartments consistent with PVS structures^19,21^. In particular, high-resolution intrathecal MRI has identified circumferential contrast enhancement surrounding major cerebral arteries, consistent with periarterial compartments accessible to CSF transport^20^. However, the observed signals remain largely restricted to CSF-accessible compartments and major surface arteries, and are inherently anatomically non-specific. Current human MRI approaches cannot resolve the fine geometry or network-level organization of PVS *in vivo*, nor can they distinguish contributions from vascular, subarachnoid, and interstitial compartments at the level of individual vessels.

Similar limitation applies to *ex vivo* high-resolution MRI studies. While post-mortem imaging following *in vivo* tracer infusion can reveal vessel-associated and ring-like enhancement patterns suggestive of a connected perivascular network^30,31^. These measurements are performed outside physiological conditions and are subject to tracer redistribution, loss of vascular dynamics, and partial-volume effects. Consequently, despite high nominal spatial resolution, these approaches do not provide a definitive *in vivo* framework for resolving the structure and organization of PVS. Together, existing MRI approaches capture brain-wide tracer transport or macroscopic perivascular features, but lack the ability to resolve a vessel-anchored, brain-wide perivascular network *in vivo*.

### Vessel-Resolved In Vivo Mapping of PVS Using Dual-Contrast MRI

To overcome these limitations, we developed an ultra-high-resolution dual-contrast MRI framework that enables vessel-resolved mapping of PVS *in vivo* across the whole brain. This capability arises from the combination of ultra-high-field MRI, implantable RF coil for enhanced sensitivity and spatial specificity^32^, and Gd-enhanced T1 contrast, which together provide sufficient signal- and contrast-to-noise ratios to resolve perivascular signal at the current spatial scale.

Using this approach, we provided *in vivo* evidence that PVS are hierarchically organized along vascular trees and extend across both surface and deep brain regions. Prominent Gd enhancement observed along the MCA and ACA branches (Fig. 1 and Supplementary Movie 1,5), including characteristic branching geometries such as T-shaped junctions, revealed that PVS follow vascular topology rather than appearing as isolated compartments. At the level of penetrating vessels, quantitative analysis showed that ∼6% of vessels exhibit detectable PVS enhancement (Fig. 2), to our knowledge, providing the first vessel-resolved estimate of PVS prevalence across the cortical vasculature. This estimate reflected detectable PVS under the current resolution and contrast conditions, which may underestimate the true prevalence of microscopic PVS. Meanwhile, the vessel diameters associated with detectable PVS were not significantly different from those without enhancement, suggesting that PVS visibility was not determined by vessel caliber or simple flow-related factors, but may instead reflected heterogeneous organization across specific vascular trees. This raises the possibility that PVS were selectively associated with particular vascular pathways, a feature that has not been resolved in prior MRI studies and warrants further investigation.

At the whole-brain level, PVS were observed not only along intracortical vessels but also across subcortical structures, including the hippocampus, midbrain, basal ganglia, and hypothalamic regions (Supplementary Movies 3-5), where they exhibited complex and spatially extended organization. As shown in Fig 4, Gd-enhanced signal was not always confined to a thin annular compartment immediately surrounding the vessel lumen, but could extend into adjacent fluid-filled compartments that remain spatially distinct from vascular signal in the hippocampus, consistent with CSF-accessible spaces rather than blood-derived contrast. This distinction is critical, as prior MRI studies have highlighted the challenge of separating perivascular signal from vascular or inflow-related contributions, and have implemented specific controls to address these confounds^22^. In our data, the absence of co-localized susceptibility distortion in phase maps further argues against vascular or susceptibility-related origins (Supplementary Fig. 2). Together, these observations demonstrated the ability of ultra-high-resolution MRI to capture brain-wide perivascular architecture that bridges vascular hierarchies and anatomical domains.

A central question raised by these observations is how PVS structures, typically only a few microns wide in optical and histological studies, can be detected at ∼30 μm MRI resolution. We do not interpret this as direct visualization of the microscopic PVS boundary. Instead, the observed signal likely reflected a voxel-scale representation of tracer-enriched fluid associated with PVS. The use of ultrashort TE, short TR, and relatively large flip angle enhanced sensitivity to Gd-driven T1 shortening while minimizing iron-induced extravascular susceptibility contamination. In addition, during the prolonged 3D acquisition, Gd was likely sampled not only within the immediate perivascular compartment but also within adjacent fluid-accessible spaces and nearby parenchyma, allowing vessel-associated enhancement to emerge through partial-volume effects. Thus, the detectability of PVS in the present study arises from the combination of strong T1 weighting, reduced susceptibility bias, and spatially organized tracer distribution near vessels, rather than from direct microscopic resolution of a 2-3 μm anatomical space.

### Implications for PVS-Mediated CSF Transport In Vivo

Although the mechanistic basis of glymphatic transport remains under active debate^9,10,33^, the present work provided an anatomical imaging framework for localizing perivascular tracer dynamics across vascular hierarchies *in vivo*. Following ventricular infusion, Gd contrast rapidly spread from the lateral ventricle into the hippocampal region, with vessel-resolved PVS enhancement detectable by 15 min (Fig. 4, Supplementary Movie 2). These findings demonstrated that tracers introduced into ventricular CSF could access a broadly distributed vessel-associated network, extending into deep cortical and subcortical regions rather than remaining confined to surface pathways^23,25^.

This global, vessel-associated tracer accessibility is consistent with prior models of CSF-ISF exchange, which propose that transport arises from a combination of periarterial pathways, interstitial diffusion, and low-velocity convective components^5,9,10,23,25,31,34-36^. Effort to identify the driving forces underlying glymphatic transport suggests that tracer movements with PVS may be influenced by arteria pulsation and or a small net forward component of CSF flow^9,37,38^. However, theoretical and fluid dynamic analyses indicate that physiological pulsations alone are insufficient to sustain long-range directional transport^35^. Instead, solute transport in the brain is more consistently described as a coupled advection-diffusion process shaped by pathway geometry, hydraulic resistance, and exchange between perivascular and interstitial compartments^39,40^.

Consistent with this view, our results demonstrate spatially organized, vessel-selective tracer transport across brain-wide perivascular structures, with only a subset of vessels (∼6%) exhibiting detectable PVS enhancement. Also, the use of a relatively low infusion volume and rate (1-2 μl over 10-20 min; 100nl/min) reduces the likelihood that the observed global PVS enhancement is dominated by infusion-driven perturbations. Nonetheless, even low-volume ventricular infusion can alter intracranial pressure and influence pathway recruitment, and tracer distribution is known to be sensitive to infusion conditions and physiological state^9,34,35,37,41^. Accordingly, our findings are best interpreted not as a direct measurement of unperturbed physiological CSF flow, but as evidence that PVS-associated tracer transport can be detected globally and anatomically localized *in vivo* under relatively mild perturbation conditions.

### Limitations and technical considerations

Several technical considerations should be noted when interpreting the present findings. First, the use of intravascular iron contrast introduces local susceptibility effects that influence both vascular and perivascular signal measurements. While iron-induced T2* shortening effectively suppresses intravascular inflow signals, facilitating separation of vessel lumens from surrounding compartments, it also generates local magnetic field perturbations that extend into the extravascular space. These susceptibility effects may partially attenuate Gd-enhanced perivascular signals and reduce local CNR. Although the use of short echo times and ultra-high spatial resolution mitigates these effects, residual susceptibility-induced bias is likely to persist, potentially leading to an underestimation of detectable PVS across the vascular population. Second, although multiple lines of evidence support that the observed perivascular enhancement reflects true tracer accumulation rather than susceptibility-related artifacts, complete exclusion of all confounding effects remains challenging. Several observations argue against a susceptibility-driven origin. The spatial organization of perivascular signal along vascular trees, including characteristic branching patterns such as T-shaped structures along ACA branches or penetrating vessel branches in deep brain regions, is not consistent with local field distortion. In addition, the detection of PVS is independent of vessel size, further arguing against susceptibility-driven contrast, which would scale with vessel caliber. Finally, perivascular enhancement is abolished following Gd washout at ∼3 h, whereas iron-induced vascular hypointensity persists, indicating dissociation between susceptibility effects and perivascular signal. Together, these findings support a tracer-based origin of the observed PVS signals, although subtle interactions between susceptibility and T1-weighted contrast cannot be entirely excluded. Third, the present framework primarily provides a structural and quasi-static mapping of perivascular tracer distribution, rather than a direct measurement of dynamic CSF flow within PVS. Capturing the spatiotemporal dynamics of tracer transport at the level of individual vessels would require substantially higher temporal resolution while maintaining mesoscale spatial resolution, which remains technically challenging. Although Gd-induced T1 shortening enhances sensitivity to CSF-associated signal, achieving sufficient CNR for resolving low-velocity flow within narrow perivascular compartments remains difficult. Future developments combining accelerated acquisition strategies with vessel-targeted analysis may enable slice-selective or vessel-resolved measurements of CSF transport along defined vascular trajectories.

Together, these findings establish a previously unavailable anatomical framework for resolving perivascular tracer dynamics across the brain *in vivo*. By enabling vessel-resolved mapping of PVS across vascular hierarchies, this approach bridges global tracer imaging and mechanistic models of CSF transport. This framework provides a foundation for future studies aimed at disentangling the relative contributions of pressure-driven flow, pulsation, diffusion, and pathway-specific resistance in brain fluid transport.

## Methods

### Animals

Adult C57BL/6 mice (4-6 months old; n = 16, 8 males and 8 females) were used in this study. Not all animals were included in every experiment due to region-specific experimental targeting of whole brain versus hippocampal structures. Subsets of animals were analyzed for each experiment, as indicated in the figure legends. Animals used exclusively for RF-coil optimization were not included in the reported totals, as the procedures were part of previously published work. Animals were group-housed (3-4 per cage) under a 12-h light/dark cycle with ad libitum access to food and water. All procedures were approved by the Massachusetts General Hospital Institutional Animal Care and Use Committee (IACUC) and complied with the National Research Council’s *Guide for the Care and Use of Laboratory Animals*.

*Sample size, replication, and inclusion/exclusion criteria*: No statistical methods were used to predetermine sample size. Sample sizes were comparable to those commonly used in ultra-high-field preclinical fMRI studies. Datasets were excluded prior to analysis if animals did not survive scanning, or suboptimal coil placement.

### Animal Surgery

#### RF coil implantation

Mice underwent surgery to affix a custom-built RF coil to the skull. Anesthesia was induced with 5% isoflurane delivered in medical air supplemented with O_2_(15-20%) for 2-3 min, and maintained at 1.5-2.0% isoflurane with respiratory rate used to monitor anesthetic depth. Ophthalmic ointment was applied to protect the corneas, and body temperature was maintained at 37±1 °C using a thermostatically controlled warming pad throughout the procedure. Animals were placed in a stereotaxic frame and secured with ear bars and a bite bar to ensure stable head positioning. Ethiqa XR (buprenorphine extended-release), 3.25 mg/ kg, s.c., was administered once pre-operatively; per protocol, no additional post-operative analgesia was required due to its prolonged effect.

The scalp was shaved and sterilized with alternating ethanol and iodine swabs. A midline incision was made to expose the skull over the region corresponding to the RF-coil footprint. Residual soft tissue was removed and the skull surface was cleaned sequentially with 0.3% H_2_O_2_and PBS, then allowed to dry. The RF coil was positioned directly over the skull above the targeted brain region, and its tuning was verified at the ^1^H Larmor frequency according to 14 T. The coil was temporarily held in place while a thin layer of cyanoacrylate adhesive was applied to bond it to the skull; curing required approximately 5–8 min. Two-part dental cement (Stoelting Co., Wood Dale, IL) was then mixed and applied to encapsulate the coil and exposed bone, taking care to secure the base of the coil firmly and prevent cement from flowing toward the eyes. Craniotomies were made above the right lateral ventricle (LV) under the microscope. 100nm/g Gd-filled capillary tube (Silica, Polyimide Coated Smooth Solid Tubing, 0.24mm, Digikey, MN), pre-connected to a micro-syringe (10 µL, WPI, FL), was inserted into the LV with the following coordinates: caudal 0.59mm, lateral 1.25mm, ventral 1.85mm from bregma. After cement placement, the scalp edges were approximated and closed using tissue adhesive. Once the cement had fully hardened (∼10 min), animals were removed from the stereotaxic frame and transferred to a warmed recovery cage until fully ambulatory. After coil fixation and stabilization, mice remained under anesthesia and were immediately transported for imaging.

#### Animal preparation for MRI

For MRI experiments, mice were imaged under free-breathing medetomidine anesthesia in combination with low-dose isoflurane. Animals were initially anesthetized with isoflurane (5% induction in medical air with supplemental 15-20% O_2_). A subcutaneous bolus of medetomidine (0.05 mg/kg) was then administered, after which isoflurane was reduced and maintained at 0.5-1.0%, titrated slowly over the course of the imaging session (4-5h total duration) to maintain physiological stability. Anesthesia was maintained with a continuous low-dose medetomidine infusion (∼0.1 mg/kg/h, s.c.). Ophthalmic ointment was applied to protect the cornea, and body temperature was maintained at 37 ±1 °C with a thermostatically regulated warm-air heating system.

In a subset of animals (n = 4), MRI experiments were performed under an alternative anesthesia regimen using ketamine/xylazine (K/X) to validate the robustness of the findings across anesthetic conditions. Mice were anesthetized with an intraperitoneal injection of ketamine (80–100 mg/kg) and xylazine (5–10 mg/kg). Supplemental doses (approximately 20-30% of the initial dose) were administered as needed to maintain a stable anesthetic depth during imaging. Respiration rate and body temperature were continuously monitored, and body temperature was maintained at 37 °C using a thermostatically controlled heating system. Importantly, no qualitative or quantitative differences were observed in the measured imaging outcomes between the medetomidine– isoflurane and K/X anesthesia conditions. Therefore, data from both anesthesia protocols were considered comparable and pooled for analysis.

Animals were positioned in a custom MRI-compatible head holder incorporating ear bars and a bite bar to minimize motion while preserving natural respiration. No paralytics were used. Physiological parameters, including respiration rate and body temperature, were continuously monitored using a small-animal monitoring system throughout the procedure (SA Instruments, NY). Once stable under medetomidine/isoflurane anesthesia, animals were transported into the MRI bore for MR scanning.

### MRI methods

#### MRI systems

All experiments were performed on horizontal bore MRI scanners (Magnex Scientific, UK, 130mm bore size) at the Athinoula A. Martinos Center for Biomedical Imaging (Charlestown, MA, USA). The 14 T (600 MHz for ^1^H) system was equipped with a Bruker Avance NEO console running ParaVision 360 v3.3 and a micro-imaging gradient set (Resonance Research, Inc.) providing peak gradient strength 1.2 T/m over a 60-mm diameter.

#### Home-built implantable RF coils

Custom implantable RF coils were fabricated in-house on thin PCB substrates following a previously described PCB-based framework. Both coils shared identical construction and matching approaches, differing only in loop geometry and resonance. A single-loop circular transceiver (≈ 6 mm diameter) was tuned and matched to 50 Ω at the ^1^H Larmor frequency at 14 T (∼600 MHz). Resonance and match were set using distributed NP0/C0G capacitors on the PCB to adjust the effective loop inductance and capacitance.

### MRI data acquisition

A 3D gradient echo (GE) sequence was used with the following parameters: TR = 80 ms; TE = 2.22 ms; flip angle = 70°; matrix size = 360 × 270 × 320; field of view = 10.8 × 8.1 × 9.6 mm^3^; spatial resolution = 30 × 30 × 30 μm^3^. The total scan time was approximately 77 min. We first acquired inflow maps, which highlighted vessel voxels as vascular control maps. Subsequently, another set of images was acquired during intraventricular Gd infusion (100 nL/min; total volume 1–2 μL) and iron injection 5mg/kg of MION (5mg/kg of MION, Molday Ion, BioPAL, Inc., Worcester, MA) via tail vein 5 minutes before MRI acquisitions.

Besides mapping cumulative glymphatic transport using whole-brain imaging, we acquired high-resolution 2D images covering the hippocampus. A 2D GE sequence was used with the following parameters: TR = 120 ms; TE = 2.95 ms; flip angle = 70°; matrix size = 320 × 320; field of view =6.4 × 6.4 mm^2^; in-plane resolution = 20 × 20 μm^2^; slice thickness = 250 μm; 2 slices. The total scan time was 7 min 42 s.

For the phase difference map validation, a 3D multi-echo gradient echo (MGE) sequence was used with the following parameters: TR = 80 ms; TE1 = 2.55 ms, TE2 = 6.56 ms; flip angle = 70°; matrix size = 360 × 270 × 320; field of view = 10.8 × 8.1 × 9.6 mm^3^; spatial resolution = 30 × 30 × 30 μm^3^.

For the Gd-based dynamic contrast-enhanced (DCE) MRI, a multi-slice MGE sequence was used with the following parameters: TR = 80 ms; TE = 2.5, 5.53, 8.56, 11.60, 14.63, and 17.66 ms; flip angle = 75°; matrix size = 256 × 256; field of view = 12.8 × 12.8 mm^2^; in-plane resolution = 50 × 50 μm^2^; slice thickness = 250 μm; 3 slices; 4 averages; and 150 repetitions.

### Data analysis

All MRI data were processed using custom pipelines implemented in MATLAB (MathWorks, Natick, MA) and Analysis of Functional NeuroImages (AFNI) software (NIMH, MD). The analysis framework was designed to systematically identify and quantify perivascular space (PVS) signals relative to vascular structures across multiple spatial scales, while maintaining consistency across animals and imaging conditions. Detailed, figure-specific analysis procedures are provided in the Supplementary Methods.

Vascular structures were first delineated from inflow-sensitive anatomical images and used as spatial references for all subsequent analyses. Gd-enhanced datasets were co-registered to the vascular maps using rigid-body alignment to ensure accurate spatial correspondence between vessels and surrounding perivascular signals. To characterize PVS organization along major vessels, vascular centerlines were extracted and used to define local coordinate systems. Intensity profiles were sampled along directions orthogonal to the vessel axis, enabling quantification of radial signal distributions. Profiles were spatially aligned relative to the vessel center and aggregated across cross-sections and animals to generate population-level estimates of perivascular signal organization.

At the level of cortical penetrating vessels, localized image patches were analyzed to quantify perivascular enhancement relative to adjacent parenchyma. Vessels were segmented based on intensity contrast, and radial signal measurements were obtained from vessel cores and surrounding tissue compartments. Perivascular signal was defined based on relative intensity differences between vessel-adjacent regions and distal tissue. To assess statistical significance, empirical null distributions were generated using randomized spatial sampling, and detection thresholds were defined relative to these controls. To examine laminar organization, measurements were grouped according to cortical depth and analyzed to compare perivascular signal distributions across layers. Depth-dependent profiles were computed by averaging radial signal changes within each cortical compartment.

Temporal dynamics of PVS signal were evaluated by comparing co-registered datasets acquired at multiple time points within the same animals. Vessel-specific measurements were matched across time, enabling paired analyses of perivascular signal evolution following tracer infusion. Similar vessel-resolved segmentation analysis was applied for hippocampal PVS signal quantification. The signal differences across baseline, post-iron, and post-Gd conditions were quantified based on the vessel cores and adjacent perivascular and parenchymal compartments. This approach enabled direct comparison of contrast contributions across imaging conditions while controlling for spatial variability.

To assess whether observed perivascular signals reflected true tracer-related contrast rather than susceptibility-induced artifacts, phase-difference and B_0_ field maps were reconstructed from multi-echo data and examined at corresponding spatial locations.

Statistical analyses were performed using paired or unpaired two-tailed t-tests as appropriate. Data are reported as mean ± s.e.m. unless otherwise specified. All analysis procedures were implemented using consistent criteria across datasets without dataset-specific parameter tuning.

## Supplementary Methods

### Quantification of PVS signal along major ACA vessels

Vascular inflow images and the corresponding Gd-enhanced PVS images were co-registered and analyzed in MATLAB. For each animal, a mid-sagittal slice covering the major ACA branches was selected (Fig. 1A). The analysis proceeded in four steps: ACA centerlines were segmented from the vascular image, perpendicular sampling lines were placed along these centerlines to extract paired V and PV intensity profiles, cross-sections with clearly resolved perivascular signal were retained, and the retained profiles were aligned to the vessel center and averaged across animals to produce the population radial profiles in Fig. 1C & D.

For ACA centerline segmentation, the vascular image of the selected slice was scaled to twice the slice maximum, smoothed with a Gaussian filter (σ = 2 pixels), and contrast-adjusted with imadjust. A binary vessel mask was obtained by Otsu thresholding with the threshold empirically scaled by a factor of 1.5, and the mask was then skeletonized to a one-pixel-wide centerline. Skeleton components smaller than 10 pixels were discarded. The cleaned skeleton was converted to a pixel graph, and at each graph edge a straight line of length 20 voxels (600 μm at 30 μm voxel size) was constructed perpendicular to the local vessel orientation and centered on the edge midpoint. Paired vascular and PVS intensity profiles were then sampled along this perpendicular line on the integer image lattice, yielding one cross-section per edge. Cross-sections were retained only if they showed a clearly resolved perivascular signal, defined as a single dominant vascular peak located near the profile center and flanked on both sides by distinct PVS peaks with a local PVS minimum at the vessel center. An example of the cross-section with its perpendicular sampling line is shown for the vascular and PVS images in Fig. 1B.

For population averaging, every accepted V and PV profile was linearly resampled to a common length of 20 points on a centered spatial axis so that the vessel center was aligned across cross-sections. Resampled profiles were averaged across all accepted cross-sections, and the standard error at each position was computed as the across-cross-section standard deviation divided by the square root of the number of non-missing values at that position. Two versions of the population profile were generated, one in raw intensity units (Fig. 1C) and the other in which each individual profile was z-scored prior to resampling (Fig. 1D). Spatial position is reported in micrometers relative to the vessel center using a 30 μm voxel size.

### Quantification of PVS signal in the parenchyma

Gd-enhanced PVS images were analyzed in MATLAB to detect perivascular enhancement around individual penetrating vessels in the cortical parenchyma, to test whether this enhancement exceeded the level expected by chance, to examine how it varied with cortical depth, and to track how it evolved from the infusion time point to 3 h post-infusion on the same vessels. The analysis was carried out on 2D patches cropped from axial and coronal slices in cortical regions, and all quantifications were performed in MATLAB.

To localize penetrating vessels and extract local image patches for analysis, the 3D PVS volume of each animal was first rotated around the vertical axis to align penetrating vessels with the image grid. The in-plane center of the cortical region of interest was selected interactively on an axial slice, and the cortical depth at that location was then selected on the corresponding coronal view. Another two adjacent axial slices spaced by one voxel above and below were also extracted at the chosen center point. From each slice a 41 by 41 voxel patch (roughly 1230 by 1230 μm at 30 μm voxel size) centered on that position was cropped. For each animal, the selection was repeated three times at each cortical depth to sample multiple spatial locations within each layer. This procedure was repeated at three cortical depths corresponding to 0 to 300, 300 to 600, and 600 to 900 μm from the pial surface, reported below as the surface, middle, and bottom layers.

To segment individual penetrating vessels and their surrounding parenchymal layers, intensities within each patch were rescaled to [0, 1] and a binary vessel mask was obtained by adaptive thresholding (adaptthresh, sensitivity 0.3, dark foreground), followed by morphological closing with a disk structuring element of radius 1 voxel. The connected components smaller than 2 voxels were removed as noise. Each connected component in the resulting mask was treated as one penetrating vessel cross-section (Fig. 2B). For every vessel, three successive single-voxel dilations of the vessel mask defined three concentric ring-shaped layers at radial distances of roughly 30, 60, and 90 μm from the vessel edge, while the original component served as the vessel core. The maximum intensity within the core and within each of the three rings was extracted and labeled V, PV1, PV2, and TS, where TS was taken as the outer parenchymal reference. For each vessel, perivascular enhancement was quantified as the relative PV-to-tissue contrast, defined as (max (PV1, PV2) - TS) / TS. The distribution of this contrast across all segmented vessels was summarized as a histogram overlaid with a kernel density estimate (Fig. 2C, left).

To test whether the observed PVS enhancement exceeded the level expected under the null hypothesis of no perivascular signal, the experimental contrast distribution was compared against a control distribution obtained by randomizing vessel positions. The control distribution was generated by repeating the entire layer extraction 1000 times on the same image patches, with the vessel positions randomized on each repetition, and summarizing the pooled values as a kernel density estimate across the 1000 repetitions. The experimental and control distributions were then overlaid (Fig. 2C, right). A detection threshold was set at the 95th percentile of the control distribution, and vessels with a contrast above this threshold were considered to carry detectable perivascular enhancement. The red shaded area in the inset of Fig. 2C highlights the portion of the experimental distribution that lies above this threshold and exceeds the control distribution, representing the fraction of vessels carrying PVS signal above the chance level.

To compare perivascular enhancement across cortical depths, the same quantification was carried out separately on the surface, middle, and bottom layers. At each depth, only vessels with a contrast above the 95th-percentile detection threshold were retained. For every retained vessel, the raw intensities at the vessel core and the three concentric rings were expressed as percentage signal change relative to the core, so that all vessels started from a common baseline at radial distance zero. The resulting radial profiles were averaged across vessels within each depth and are reported as mean plus or minus s.e.m. (Fig. 2D). Differences between the three depths at the 60 and 90 μm rings were evaluated by paired t-test.

To test whether the ability to detect PVS signal was confounded by vessel caliber and therefore reflected a partial-volume effect rather than true perivascular enhancement, the number of voxels in each segmented vessel mask was taken as a proxy for vessel size, and vessel sizes were compared between detectable and undetectable vessels by two-sample t-test (Fig. 2E). Segments exceeding 20 voxels (600 µm) were excluded, as they fall outside the characteristic spatial extent of individual penetrating vessels. Because vessel size distributions were heavily right-skewed and violated the normality assumption required by parametric tests (Shapiro-Wilk test, p < 0.001 for both groups), a two-sample Kolmogorov-Smirnov test was used to assess whether the two groups shared the same underlying distribution (MATLAB kstest2). The two-sample KS test showed no significant difference in vessel size distribution between PVS-detectable (n = 90, median = 4 voxels) and PVS-undetectable vessels (n = 1444, median = 3 voxels) (p=0.3346).

To test whether the detected PVS signal reflected true perivascular Gd accumulation rather than an imaging artifact or an iron-induced susceptibility effect, two additional analyses were performed on the pooled vessel population. First, to verify that the enhancement was specific to the infusion time point and not a persistent feature of the image, the raw intensities at the vessel core and the three concentric rings were averaged across detectable vessels at infusion and at 3 h post-infusion for comparison (Fig. 3B). Second, to track how PVS enhancement evolved over time on the same vessels, the infusion and 3 h post-infusion image volumes from each animal were co-registered, and the vessel masks generated at the infusion time point were reapplied to the post-infusion volume. This ensured that the vessel core and the three concentric rings were sampled at identical spatial locations at both time points, so that each vessel contributed one paired measurement. For every vessel passing the 95th-percentile detection threshold at infusion, the infusion and post-infusion relative PV-to-tissue contrast values were compared by paired t-test (Fig. 3C, 131 vessels).

### Quantification of PVS signal around hippocampal penetrating vessels across imaging contrasts

To confirm that the perivascular Gd enhancement observed around hippocampal penetrating vessels reflected true Gd accumulation in the perivascular space rather than a feature already present at baseline or introduced by the preceding iron infusion, signal intensities were compared across three sequential scans of the same animals: a baseline scan, a post-iron infusion scan, and a post-Gd infusion scan. The three scans were co-registered so that the same vessel and the same surrounding locations could be sampled at each time point. The analysis was carried out in MATLAB on coronal slices covering the dorsal hippocampus.

For every penetrating vessel, three ROIs were placed along a radial line centered on the vessel, one on the vessel lumen, one on the perivascular space, and one on the surrounding tissue, corresponding to radial distances of 0, 20, and 40 μm from the vessel center (Fig. 4D inset). The same three ROIs were then applied to the baseline, post-iron, and post-Gd scans of that vessel, yielding one intensity value per ROI per scan. A vessel was retained only if, on the post-Gd scan, the perivascular ROI was brighter than both the lumen ROI and the tissue ROI, which left 38 vessels from 8 animals. To place the three scans on a common scale despite their different absolute intensity ranges, the three ROI values of each vessel were z-scored within that vessel and within that scan, averaged across the 38 retained vessels separately for each scan, and plotted as mean plus or minus s.e.m. as a function of radial distance from the vessel center (Fig. 4D).

### Phase-map validation of PVS signal

To test whether the bright perivascular signals detected on the MGE images reflected true Gd-induced T1 shortening in the perivascular space rather than phase-related artifacts such as local susceptibility offsets, chemical shift, or partial-volume phase cancellation, phase-difference and B0 field maps were reconstructed from the same raw multi-echo GRE k-space data used for the image-based PVS analysis and inspected at the same spatial locations as the bright perivascular voxels.

Phase-difference maps between the two echoes were computed as the angle of the Hermitian product HP = S2 · conj(S1), where S1 and S2 are the complex images at the first and second echoes. This Hermitian formulation cancels any echo-independent background phase, such as coil phase offsets, and yields a wrapped phase-difference map whose value at each voxel depends only on the local frequency offset and the echo-time difference. The wrapped phase-difference map was then unwrapped in 3D using a region-growing phase-unwrapping algorithm (https://github.com/geggo/phase-unwrap), and the unwrapped phase difference was converted to a B0 field map in Hz by B0 = −Δϕ / (2π · ΔTE), where ΔTE was the difference between the two echo times.

## Data availability

The imaging datasets generated in this study will be deposited in a public repository prior to publication and made accessible via a unique accession code. The archive will include preprocessed datasets and statistical maps sufficient to reproduce the primary analyses. Raw imaging data, including k-space datasets and associated calibration scans, will be made available through the repository.

## Code availability

Custom scripts used for MRI preprocessing, reconstruction, statistical modelling will be deposited in a public repository (github) prior to publication. Acquisition sequence components and hardware-specific implementations associated with ultra-high-field MRI systems may require coordination with the scanner vendor.

## Acknowledgements

This work was supported by the Alzheimer’s Association (ARFD-23-1145375 to X.A.Z.), the Brain & Behavior Research Foundation Young Investigator Award (to X.A.Z.), and NIH grant (R01AG094755 to X.A.Z). Additional support was provided by NIH grants (RF1NS124778, R01NS122904, U19NS123717), NSF grant 2123971, and the S10 instrumentation grants (S10MH124733, S10OD036211) to the Martinos Center.

## Figure Legends

**Supplementary Figure 1.**
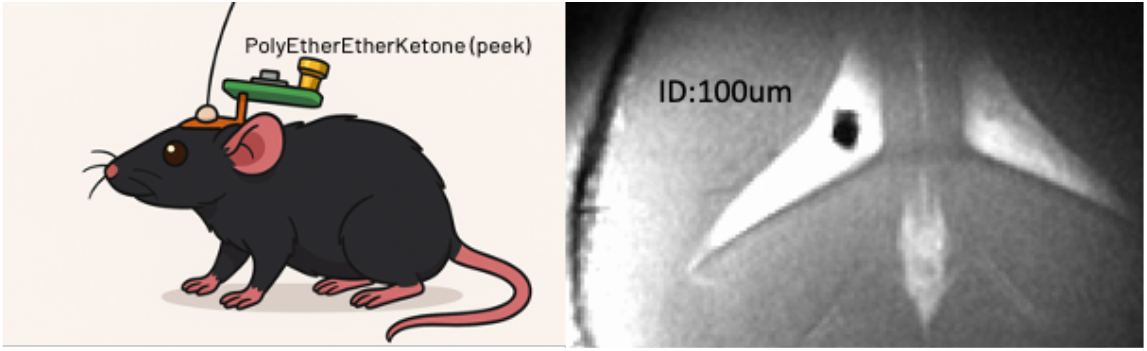
Schematic of the experimental setup. A mouse with a custom PEEK head implant for imaging (left) and the corresponding MR image showing implant geometry (right, inner diameter = 100 μm).

**Supplementary Figure 2.**
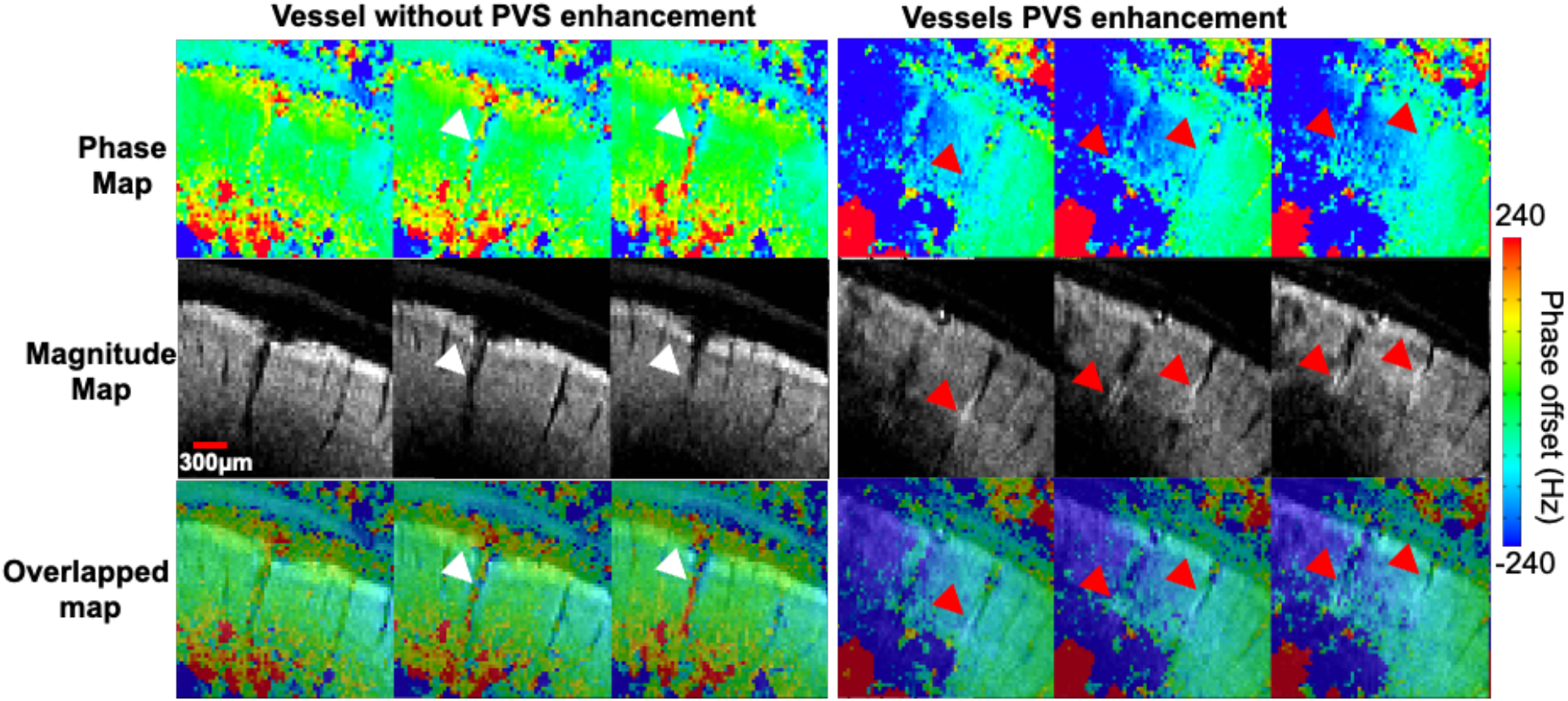
Perivascular enhancement is not driven by local susceptibility effects. Representative phase (top) and corresponding magnitude (middle) images at the level of individual cortical penetrating vessels, with overlaid maps (bottom). White arrowheads indicate vessels exhibiting strong local field offsets, reflected as pronounced phase shifts, but without corresponding perivascular signal enhancement in magnitude images. In contrast, red arrowheads indicate vessels showing robust perivascular enhancement in magnitude images, despite minimal phase perturbation. These observations demonstrate a dissociation between local susceptibility effects and perivascular signal, indicating that perivascular enhancement is not attributable to iron-induced field inhomogeneity.

**Supplementary Figure 3.**
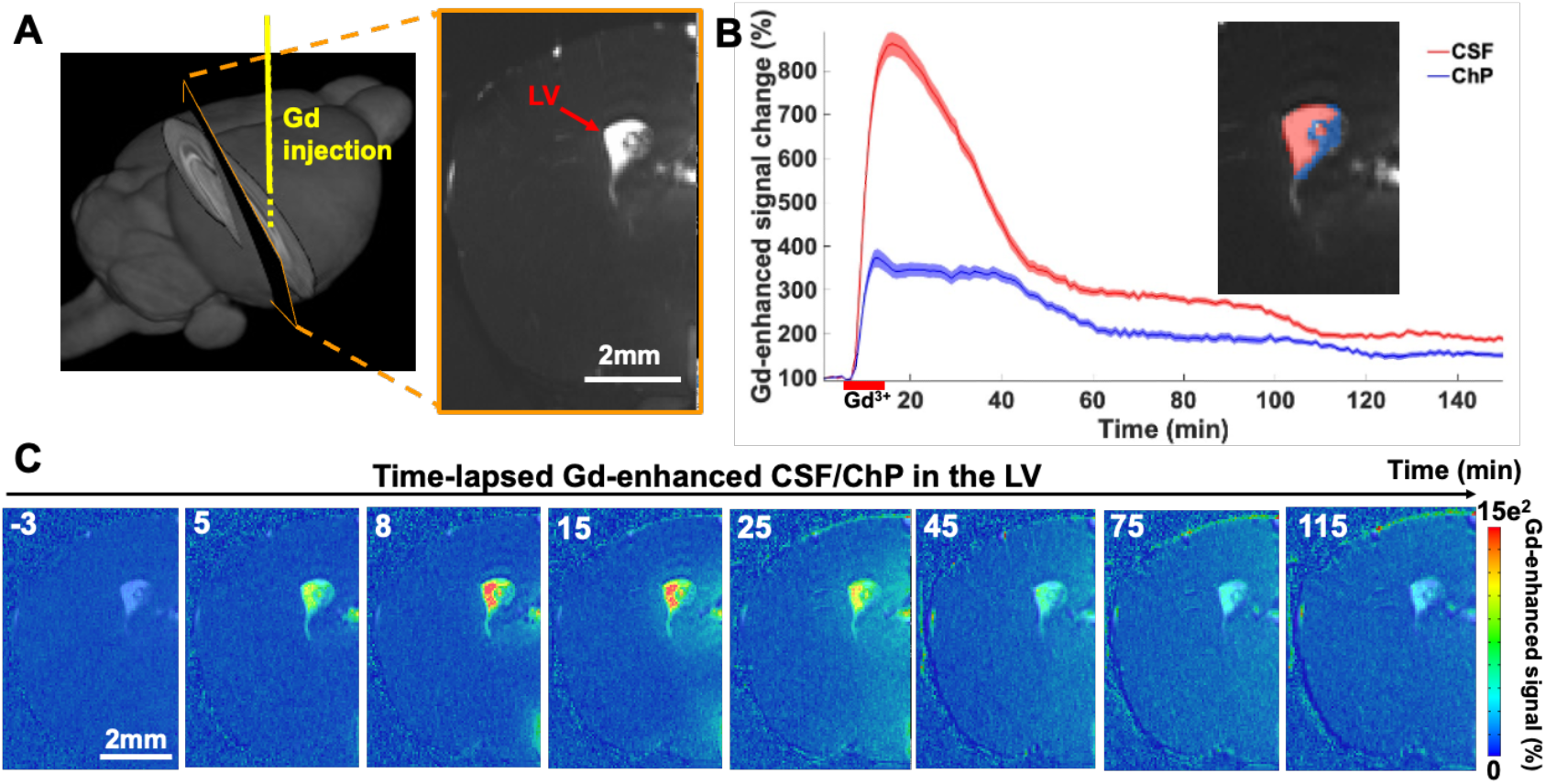
Ventricular infusion reveals dynamic tracer accumulation in CSF and choroid plexus. **A**, Schematic of intraventricular Gd infusion into the lateral ventricle (LV) with corresponding MR image showing catheter placement. **B**, Representative time courses of Gd-enhanced signal following infusion, showing rapid signal increase in CSF and delayed, lower-amplitude accumulation in the choroid plexus (ChP). Data are shown as mean ± s.e.m. across voxels within each ROI. Inset, ROIs for CSF (red) and ChP (blue). **C**, Time-lapse Gd-enhanced images of the LV following infusion, demonstrating rapid tracer spread within free-moving CSF, with peak signal occurring within ∼8–10 min and minimal accumulation in ChP-associated regions.

**Supplementary Movie 1. Slice-by-slice visualization of the perivascular space surrounding the middle cerebral artery**.

Sequential MRI slices demonstrate the CSF-filled pial perivascular space associated with the middle cerebral artery (MCA), illustrating its spatial continuity along the vessel trajectory.

**Supplementary Movie 2. Dynamic Gd-enhanced MRI mapping following lateral ventricular infusion**.

Time-resolved MGE images show the spatiotemporal progression of Gd-enhanced signal following lateral ventricular infusion. Signal time courses are shown for three ROIs: LV-CSF in red, hippocampal CA1 in yellow, and hippocampal CA3 in green. Images were acquired at 50 × 50 μm in-plane resolution with a temporal resolution of approximately 56 s per frame.

**Supplementary Movie 3. Slice-by-slice visualization of perivascular spaces surrounding vessels supplying the hippocampus and basal ganglia**.

Sequential MRI slices demonstrate vessel-associated PVS along vascular pathways supplying the hippocampus and basal ganglia, illustrating their spatial organization across adjacent imaging planes.

**Supplementary Movie 4. Slice-by-slice visualization of perivascular spaces surrounding vessels supplying the dorsalmidbrain**.

Sequential MRI slices demonstrate vessel-associated PVS along vascular pathways supplying the dorsal midbrain, illustrating their spatial organization across adjacent imaging planes.

**Supplementary Movie 5. Slice-by-slice visualization of perivascular spaces surrounding vessels supplying ACA branches and thalamic/hypothalamic regions**.

Sequential MRI slices demonstrate vessel-associated PVS along vascular pathways supplying the ACA branches, highlighting their spatial organization across the characteristic T-shaped branching structure (upper-right box). Additional slice-by-slice views show PVS along vessels supplying the thalamic and hypothalamic regions, including branches arising from the circle of Willis (lower-left box).

## References

1 Virchow, R. Ueber die Erweiterung kleinerer Gefäfse. Archiv für pathologische Anatomie und Physiologie und für klinische Medicin 3, 427–462 (1851). 10.1007/BF01960918

2 Robin, C. Recherches sur quelques particularites de la structure des capillaires de l’encephale. J Physiol Homme Animaux 2, 537 (1859).

3 Ineichen, B. V. et al. Perivascular spaces and their role in neuroinflammation. Neuron 110, 3566–3581 (2022). 10.1016/j.neuron.2022.10.024

4 Iliff, J. J. et al. A Paravascular Pathway Facilitates CSF Flow Through the Brain Parenchyma and the Clearance of Interstitial Solutes, Including Amyloid β. Science Translational Medicine 4, 147ra111–147ra111 (2012). 10.1126/scitranslmed.3003748

5 Carare, R. O. et al. Solutes, but not cells, drain from the brain parenchyma along basement membranes of capillaries and arteries: significance for cerebral amyloid angiopathy and neuroimmunology. Neuropathol Appl Neurobiol 34, 131–144 (2008). 10.1111/j.1365-2990.2007.00926.x

6 Carare, R. O. Intramural Peri Arterial Drainage (IPAD) of peptides from the brain parenchyma and retina. Alzheimer’s & Dementia 21, e100903 (2025). 10.1002/alz70855_100903

7 Albargothy, N. J. et al. Convective influx/glymphatic system: tracers injected into the CSF enter and leave the brain along separate periarterial basement membrane pathways. Acta neuropathologica 136, 139–152 (2018). 10.1007/s00401-018-1862-7

8 Wardlaw, J. M. et al. Perivascular spaces in the brain: anatomy, physiology and pathology. Nat Rev Neurol 16, 137–153 (2020). 10.1038/s41582-020-0312-z

9 Mestre, H., Mori, Y. & Nedergaard, M. The Brain’s Glymphatic System: Current Controversies. Trends in neurosciences 43, 458–466 (2020). 10.1016/j.tins.2020.04.003

10 Hladky, S. B. & Barrand, M. A. The glymphatic hypothesis: the theory and the evidence. Fluids Barriers CNS 19, 9 (2022). 10.1186/s12987-021-00282-z

11 Hablitz, L. M. et al. Circadian control of brain glymphatic and lymphatic fluid flow. Nature communications 11, 4411 (2020). 10.1038/s41467-020-18115-2

12 Fultz, N. E. et al. Coupled electrophysiological, hemodynamic, and cerebrospinal fluid oscillations in human sleep. Science 366, 628–631 (2019). 10.1126/science.aax5440

13 Jiang-Xie, L.-F. et al. Neuronal dynamics direct cerebrospinal fluid perfusion and brain clearance. Nature 627, 157–164 (2024). 10.1038/s41586-024-07108-6

14 Holstein-Rønsbo, S. et al. Glymphatic influx and clearance are accelerated by neurovascular coupling. Nature neuroscience 26, 1042–1053 (2023). 10.1038/s41593-023-01327-2

15 Mestre, H. et al. Flow of cerebrospinal fluid is driven by arterial pulsations and is reduced in hypertension. Nature communications 9, 4878 (2018). 10.1038/s41467-018-07318-3

16 Hauglund, N. L. et al. Norepinephrine-mediated slow vasomotion drives glymphatic clearance during sleep. Cell 188, 606–622.e617 (2025). 10.1016/j.cell.2024.11.027

17 Chuang, K.-H. et al. Cholinergic basal forebrain neurons regulate vascular dynamics and cerebrospinal fluid flux. Nature communications 16, 5343 (2025). 10.1038/s41467-025-60812-3

18 Sun, Q. et al. Enhancing glymphatic fluid transport by pan-adrenergic inhibition suppresses epileptogenesis in male mice. Nature communications 15, 9600 (2024). 10.1038/s41467-024-53430-y

19 Naganawa, S., Nakane, T., Kawai, H. & Taoka, T. Gd-based Contrast Enhancement of the Perivascular Spaces in the Basal Ganglia. Magn Reson Med Sci 16, 61–65 (2017). 10.2463/mrms.mp.2016-0039

20 Eide, P. K. & Ringstad, G. Functional analysis of the human perivascular subarachnoid space. Nature communications 15, 2001 (2024). 10.1038/s41467-024-46329-1

21 Yamamoto, E. A. et al. The perivascular space is a conduit for cerebrospinal fluid flow in humans: A proof-of-principle report. Proceedings of the National Academy of Sciences 121, e2407246121 (2024). 10.1073/pnas.2407246121

22 Monte, B. et al. Characterization of perivascular space pathology in a rat model of cerebral small vessel disease by in vivo magnetic resonance imaging. Journal of Cerebral Blood Flow & Metabolism 42, 1813–1826 (2022). 10.1177/0271678X221105668

23 Iliff, J. J. et al. Brain-wide pathway for waste clearance captured by contrast-enhanced MRI. The Journal of clinical investigation 123, 1299–1309 (2013). 10.1172/JCI67677

24 Lee, H. et al. Quantitative Gd-DOTA uptake from cerebrospinal fluid into rat brain using 3D VFA-SPGR at 9.4T. Magnetic resonance in medicine : official journal of the Society of Magnetic Resonance in Medicine / Society of Magnetic Resonance in Medicine 79, 1568–1578 (2018). 10.1002/mrm.26779

25 Stanton, E. H. et al. Mapping of CSF transport using high spatiotemporal resolution dynamic contrast-enhanced MRI in mice: Effect of anesthesia. Magnet Reson Med 85, 3326–3342 (2021). 10.1002/mrm.28645

26 Chen, X. et al. Mapping optogenetically-driven single-vessel fMRI with concurrent neuronal calcium recordings in the rat hippocampus. Nature communications 10, 5239 (2019). 10.1038/s41467-019-12850-x

27 Mortensen, K. N. et al. Perivascular cerebrospinal fluid inflow matches interstitial fluid efflux in anesthetized rats. iScience 28, 112323 (2025). 10.1016/j.isci.2025.112323

28 Chen, X. et al. Mapping optogenetically-driven single-vessel fMRI with concurrent neuronal calcium recordings in the rat hippocampus. Nature communications 10, 5239 (2019). 10.1038/s41467-019-12850-x

29 Watts, R., Steinklein, J. M., Waldman, L., Zhou, X. & Filippi, C. G. Measuring Glymphatic Flow in Man Using Quantitative Contrast-Enhanced MRI. AJNR. American journal of neuroradiology 40, 648–651 (2019). 10.3174/ajnr.A5931

30 Magdoom, K. N. et al. MRI of Whole Rat Brain Perivascular Network Reveals Role for Ventricles in Brain Waste Clearance. Scientific reports 9, 11480 (2019). 10.1038/s41598-019-44938-1

31 Rey, J. A. et al. Perivascular network segmentations derived from high-field MRI and their implications for perivascular and parenchymal mass transport in the rat brain. Scientific reports 13, 9205 (2023). 10.1038/s41598-023-34850-0

32 Hike, D. et al. High-resolution awake mouse fMRI at 14 tesla. eLife 13, RP95528 (2025). 10.7554/eLife.95528

33 Zhao, L., Tannenbaum, A., Bakker, E. & Benveniste, H. Physiology of Glymphatic Solute Transport and Waste Clearance from the Brain. Physiology (Bethesda) 37, 0 (2022). 10.1152/physiol.00015.2022

34 Smith, A. J., Yao, X., Dix, J. A., Jin, B.-J. & Verkman, A. S. Test of the ‘glymphatic’ hypothesis demonstrates diffusive and aquaporin-4-independent solute transport in rodent brain parenchyma. eLife 6, e27679 (2017). 10.7554/eLife.27679

35 Kedarasetti, R. T., Drew, P. J. & Costanzo, F. Arterial pulsations drive oscillatory flow of CSF but not directional pumping. Scientific reports 10, 10102 (2020). 10.1038/s41598-020-66887-w

36 Morris, A. W. et al. Vascular basement membranes as pathways for the passage of fluid into and out of the brain. Acta neuropathologica 131, 725–736 (2016). 10.1007/s00401-016-1555-z

37 Bedussi, B., Almasian, M., de Vos, J., VanBavel, E. & Bakker, E. N. Paravascular spaces at the brain surface: Low resistance pathways for cerebrospinal fluid flow. Journal of cerebral blood flow and metabolism : official journal of the International Society of Cerebral Blood Flow and Metabolism 38, 719–726 (2018). 10.1177/0271678x17737984

38 Kiviniemi, V. et al. Ultra-fast magnetic resonance encephalography of physiological brain activity – Glymphatic pulsation mechanisms? Journal of Cerebral Blood Flow & Metabolism 36, 1033–1045 (2016). 10.1177/0271678x15622047

39 Ray, L. A. & Heys, J. J. Fluid Flow and Mass Transport in Brain Tissue. Fluids 4, 196 (2019).

40 Ray, L., Iliff, J. J. & Heys, J. J. Analysis of convective and diffusive transport in the brain interstitium. Fluids and Barriers of the CNS 16, 6 (2019). 10.1186/s12987-019-0126-9

41 Raghunandan, A. et al. Bulk flow of cerebrospinal fluid observed in periarterial spaces is not an artifact of injection. Elife 10 (2021). 10.7554/eLife.65958

